# Representational geometry as a fidelity metric for connectome-constrained networks: evidence from the *Drosophila* visual system

**DOI:** 10.64898/2026.06.10.731214

**Authors:** Michael G. Zhou, Jennifer O. Hasler

## Abstract

What does biological wiring actually contribute to neural computation? Behavioral experiments can test whether a model produces the right outputs, but they cannot determine whether its internal representations are biologically faithful. Brunton et al. (2026) made this concrete: a *C. elegans* worm connectome trained with deep reinforcement learning produces realistic *Drosophila* fly walking, yet the model is biologically meaningless, because behavioral fidelity is achievable without biological fidelity. We need a population-level metric that discriminates real biological wiring from arbitrary wiring, without requiring a behavioral decoder. We propose representational geometry as that metric, and test it on the Flyvis pretrained *Drosophila* visual system ensemble (Lappalainen et al., 2024) against three families of random wiring: a weight-shuffled baseline, and two topology-randomizing null schemes, degree-preserving swap and Erdős–Rényi, used through-out the rest of the paper. Using individual-pairwise RDM comparison rather than the weaker ensemble-mean statistic (which can itself exceed its own within-real ceiling, a result that is not possible under an honest comparison), real wiring is clearly distinguishable from random wiring before any training, across all three stimulus conditions tested and both topology-randomizing schemes (*p* < 10^−13^ in every case). An apparent exception on the combined ON+OFF stimulus set turns out to be a confound: it measures a coarse, shared ON/OFF-discrimination signal any network reproduces, and disappears once within-polarity structure is isolated. Grounding this in measured biology required first diagnosing and repairing the reference itself. A biological reference constructed from idealized tuning curves fit to Maisak et al. (Maisak et al., 2013) proved 97.8% explained by circular stimulus structure alone and uninterpretable as a fidelity instrument. We replaced it with a reference built from real per-cell recordings (Henning et al., 2022), substantially less circular (64–82%, depending on reconstruction method) and independently validated as carrying genuine, replicable signal via a synthetic-noise floor and split-half reliability test.

With a validated reference in hand, we tested the case Brunton’s finding makes urgent directly: does geometry distinguish real, trained wiring from random wiring that has *also* been trained to task adequacy? Two independent methods largely agree it does not. A biology-mediated comparison finds real and trained-random wiring statistically indistinguishable, once properly size-matched, for both null schemes. A direct RDM comparison requiring no biological reference at all finds the same substantial narrowing, and in one specific case, trained Erdős–Rényi on OFF-polarity structure, closes the gap completely, the only case across either method where that happens outright. Checkpoint-level trajectories show this convergence is abrupt, not gradual, and its magnitude is set entirely by how much distinguishability existed before training began, not by wiring scheme or a separate developmental process.

A negative-trending pattern in one comparison, random wiring correlating *more* strongly with biology than real wiring, motivated two further tests. First, this pattern tracks training generally, not wiring identity: a matched real-versus-untrained-random comparison is itself significant, and the finding generalizes robustly to the trained null schemes once fairly resampled. Second, retraining eight seed networks from identical wiring with different training-noise seeds, following the methodology of Frankle et al. (2020), finds no significant evidence that wiring realization, rather than training-process randomness, determines the direction an individual network’s fidelity trend takes, on the more trustworthy of our two biological references. Two networks show internal dynamics beyond simple clustering or scatter, one with a trajectory reversal mid-training that produces three qualitatively different retrained outcomes from identical wiring, suggestive of a genuine dynamical bifurcation rather than noise.

Together these results show that representational geometry is a genuine, population-level fidelity signal: it reliably distinguishes real from random wiring before training, corroborated independently by CKA, but that distinction narrows substantially and in specific cases disappears entirely once both are trained to task adequacy, the direct test Brunton’s finding raises. We report this as an honest, largely null result on the training question, not the confirmation we set out to find, together with a validated, non-circular biological reference and a diagnosed instance of the ensemble-mean statistical artifact, as resources for evaluating fidelity in connectome-scale emulations more broadly.

## 1 Introduction

A central question in systems neuroscience is whether the specific wiring of a biological circuit matters for neural computation, or whether any sufficiently capable network could produce equivalent representations and behavior. This question has become practically urgent. Complete wiring diagrams (connectomes) of the *Drosophila melanogaster* nervous system are now available at synaptic resolution (Dorkenwald et al., 2024; Schlegel et al., 2024), and computational frameworks now enable networks whose architecture is fully constrained by these connectomes (Lappalainen et al., 2024; Pugliese et al., 2025). Similar efforts are underway for mouse visual cortex (MICrONS Consortium et al., 2025). As connectome-constrained emulations grow in scale and biological detail, a principled framework for evaluating their fidelity becomes essential. Recent work has begun to survey the landscape of possible fidelity tests for whole-brain emulation at a conceptual level (Linssen and Koene, 2025), but no specific candidate metric has yet been proposed and empirically validated against connectome-constrained models.

The problem is that behavioral fidelity alone is insufficient. Brunton et al. (2026) demonstrated this sharply by constructing a virtual chimera: a *C. elegans* worm connectome controlling a *Drosophila* fly body, trained with deep reinforcement learning to imitate real fly walking kinematics. The chimera succeeded — yet the model is biologically meaningless. Its worm connectome functions simply as a recurrent neural network with rich enough dynamics to support locomotor learning, a role that a randomly connected network could fill equally well. The conclusion is direct: behavioral fidelity is achievable without biological fidelity, and behavioral evaluation cannot distinguish biological wiring from arbitrary wiring. The authors explicitly identify, but do not test, the sharper version of this problem: they never trained the connectome’s own synaptic weights, predicting that doing so would only deepen the interpretability problem their untrained-random result already raises. That prediction is the specific gap this paper closes.

What, then, should we measure instead? We propose that the answer lies at the level of population codes. Representational geometry — the structure of pairwise distances between a population’s responses to different stimuli — captures how a circuit organizes its representational space, independently of any downstream behavioral decoder. Representational similarity analysis (RSA; Kriegeskorte et al. 2008) provides a principled framework for comparing these geometries across brains, models, and species, and has been used to align neural data with model representations in the primate visual system (Yamins and DiCarlo, 2016). Subsequent theoretical work showed that representational geometry captures aspects of population coding that scalar metrics — including Fisher information — cannot (Kriegeskorte and Wei, 2021). If connectome-constrained networks produce a representational geometry that random networks do not reproduce, then geometry constitutes a candidate fidelity signal operating at the population level without requiring behavioral output.

Here we test this hypothesis using the Flyvis pretrained *Drosophila* visual system ensemble (Lappalainen et al., 2024): 50 networks whose synaptic architecture is fixed to the Flyvis connectome, trained to perform optic-flow estimation on naturalistic video. We compare these connectome-constrained (CC) networks against three families of random wiring: a weight-shuffled baseline (topology intact, weight values scrambled), and two schemes that randomize topology directly, degree-preserving swap and Erdős–Rényi, the two null schemes the trained-wiring comparison actually depends on. We measure whether these populations occupy geometrically distinct regions of population response space using RSA and CKA (Kornblith et al., 2019), and whether CC geometry is more consistent than random wiring’s with real T4/T5 direction-tuning recordings, first from published summary parameters (Maisak et al., 2013), and, once that reference proved unusable, from real per-cell data (Henning et al., 2022).

Our results support the untrained claim cleanly, once the correct statistic is used, and give a qualified, largely negative answer to the trained-wiring question Brunton’s finding raises directly. Ensemble-mean RDM correlation, averaging every model’s response pattern before comparing, understates and in one condition actively inverts the true gap between real and random wiring; individual-pairwise comparison resolves this, and the resulting untrained-distinguishability result holds for all three random-wiring families, across every stimulus condition tested, and is independently corroborated by CKA. Grounding this in measured biology required repairing the reference first: the original T4/T5 reference proved 97.8% explained by circular stimulus structure and uninterpretable, a validated replacement built from real per-cell recordings (Henning et al., 2022) is substantially, though not entirely, non-circular. Testing the trained-wiring question directly, using two independent methods, we find real and trained-random wiring are largely statistically indistinguishable, with one specific, complete exception (trained Erdős–Rényi on OFF-polarity structure) and one broader, honest null (a biology-mediated comparison that does not survive proper size-matching). A further pair of experiments traces a negative-trending pattern in one comparison to training itself rather than wiring identity, and finds no significant evidence that an individual network’s own fidelity-trend direction is determined by its wiring realization rather than training-process randomness, though two networks’ internal dynamics suggest a genuine dynamical bifurcation rather than simple noise.

For experimentalists, this framework yields a testable prediction: connectome perturbations that preserve behavior but disrupt wiring specificity should reduce geometric fidelity before behavioral deficits emerge, at least in the untrained regime where that fidelity signal is unambiguous. For theoreticians, the finding that trained real and trained random wiring converge so substantially, and in one case completely, sharpens rather than resolves the question Brunton’s finding raises: task optimization alone appears to account for most, and in specific cases all, of what looked like a wiring-specific fidelity signal before training. Together, these results suggest that representational geometry is a genuine, decoder-free fidelity signal for untrained or partially-trained emulations, but that its power to discriminate real from arbitrary wiring erodes, in places entirely, once both are optimized to the same task.

## 2 Methods

### 2.1 Neural network models

We used the pretrained Flyvis ensemble (Lappalainen et al., 2024) — 50 connectome-constrained network models of the *Drosophila melanogaster* visual system trained to perform optic-flow estimation. Architecture is fixed to the reconstructed fly connectome (retina, lamina, medulla, lobula, and lobula plate; 65 distinct cell types), leaving synaptic weights and cellular biophysical parameters as the free, trainable quantities. Each unit is a graded-potential, non-spiking, single-compartment model with first-order voltage dynamics (a leaky RC-style update toward an input-driven equilibrium; PPNeuronIGRSynapses, ReLU activation): there is no spike threshold, reset, or refractory period. Models are indexed 000–049 within the flow/0000 ensemble; all 50 were used in every experiment.

#### Training objective

A dedicated decoder (DecoderGAVP, output shape [8, 2], kernel size 5) reads the network’s pooled activity and is trained, jointly with the network’s own weights, against an L2-norm optic-flow loss plus an activity penalty, using Adam under a stepwise learning-rate schedule (5 × 10^−5^ decaying to 5 × 10^−6^) for 250,000 iterations at batch size 4. Training uses Flyvis’s own built-in training entry point rather than a custom loop, so nothing about the recipe diverges from how the pretrained ensemble itself was produced. This identical recipe, varying only which connectome constrains the network’s architecture, is what makes the real-versus-null wiring comparisons in Experiments 3–5 a controlled, single-variable test.

### 2.2 Stimuli

Three stimulus sets were used, matching the coverage of the biological reference available for each comparison:

- **ON-only:** 12 directions at 30^°^ increments (0^°^ through 330^°^), ON polarity only. Yields a 12 × 12 stimulus RDM.
- **ON+OFF:** 24 conditions — 12 directions × 2 polarities, interleaved by direction (OFF at even indices, ON at odd indices). Yields a 24 × 24 stimulus RDM.
- **Henning 8-direction:** 8 directions at 45^°^ increments, matching the native resolution of the Henning et al. (Henning et al., 2022) biological reference (Biological reference RDM, below). Yields an 8 × 8 stimulus RDM.

### 2.3 Population vectors

For each model, stimulus condition, and checkpoint, we simulated the network’s response and extracted a population vector as the peak central-cell response per cell type, yielding a 65-dimensional vector per stimulus. Every representational dissimilarity matrix (RDM) in this paper is built from these simulated response vectors, never from the connectome’s wiring diagram directly: wiring, real or null, is the independent variable that differs between conditions, and representational geometry is the downstream, measured consequence of running that wiring through simulation.

### 2.4 Null wiring schemes

Three families of random wiring were used, testing distinguishability under increasingly specific structural constraints.

#### Weight-shuffled baseline

The real connectome’s topology is held fixed; all trainable weight values are shuffled in a sign-preserving manner (Dale’s law respected). This is the baseline used to establish the original, untrained real-versus-random result (Experiment 1) and is a topologically identical, weight-scrambled counterpart to the real network.

#### Degree-preserving swap

A Maslov–Sneppen double-edge swap on the real connectome’s topology: repeatedly selecting two edges *a* →*b* and *c* →*d* and rewiring them to *a* →*d* and *c* → *b*, rejecting any swap that would create a self-loop or duplicate edge. This preserves every cell type’s in- and out-degree exactly (its number of distinct connection partners) while scrambling which specific partners those are. At the swap intensity used (target 10 × the edge count in successful swaps), approximately 39% of connection pairs changed from the real connectome.

#### Erdős–Rényi

A full topological scramble: new connection pairs are drawn uniformly at random, with no degree constraint at all, the more aggressive of the two topology-randomizing schemes.

All three schemes preserve total edge count, total synapse count, and Dale’s law sign structure by construction; only the specific pattern of which cell types connect to which differs between them, and only the degree-preserving scheme additionally preserves each cell type’s number of distinct connection partners.

### 2.5 Representational dissimilarity matrices

For each model, an RDM was computed by applying cosine distance to all pairs of population vectors (Eq. 1), with a small epsilon (10^−10^) added before distance computation to guard against zero-norm vectors.

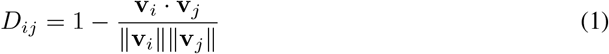

### 2.6 RSA: individual-pairwise correlation, not ensemble-mean

RDM similarity is quantified by Spearman rank correlation between the upper triangular entries of two RDMs, with significance assessed by a stimulus-label permutation test (Nili et al., 2014) (10,000 permutations, one-sided).

#### Individual-pairwise, not ensemble-mean

Averaging every model’s RDM into a single ensemble-mean RDM before correlating — computing one real-mean RDM and one null-mean RDM, then a single correlation between them — smooths over individual-model variability on both sides and can substantially understate, or in degenerate cases even invert, the true real-versus-null gap: on the ON+OFF condition, the ensemble-mean statistic (Spearman *r* = 0.846) exceeds the individual-pairwise within-real baseline itself (*r* = 0.838), a result that is not possible under an honest comparison, since real networks cannot on average be more similar to null wiring than to each other. We therefore report, as the primary statistic throughout, the individual-pairwise comparison: every real model’s RDM correlated against every other real model’s RDM (*within-real* baseline, 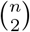 pairs) and against every null model’s RDM (*real-vs-null, n* × *n*_null_ pairs), each set of correlations compared by Mann–Whitney *U* test (within-real *>* real-vs-null). The within-real baseline serves as the realistic ceiling against which real-vs-null is judged, rather than an idealized *r* = 1.

### 2.7 Biological reference RDM

An initial biological reference constructed from idealized von Mises tuning curves fit to summary parameters in Maisak et al. (Maisak et al., 2013) was found to be dominated by circular stimulus structure rather than genuine direction-tuning signal (Results, §3.2) and is not used in this paper’s reported comparisons; we retain its construction and diagnosis here only as motivation for the replacement below.

#### Henning et al. reference

We instead constructed the biological reference from Henning, Ramos-Traslosheros, Gür & Silies (Henning et al., 2022), real per-cell T4/T5 direction-tuning recordings rather than idealized summary curves, in two independent ways:

- **Raw**. Each cell’s peak response at each of the 8 native stimulus directions is averaged directly within its cell-type group; no curve-fitting step is applied.
- **Von Mises**. Each cell’s own fitted preferred direction and concentration parameter (derived from its complex tuning-vector summary) parameterizes an individual von Mises curve; curves are averaged within each cell-type group. Unlike the retired Maisak construction, every cell contributes its own fitted direction and sharpness rather than sharing one idealized, identical-width curve per layer.

Both versions remain substantially circular (raw: 63.5% of variance explained by a pure circular-distance model; von Mises: 82.3%), meaningfully less than the retired Maisak reference (95.6%) but not negligible. Two further tests establish that the non-circular remainder is genuine signal, not reconstruction artifact:

#### Synthetic-noise floor

Pure synthetic noise, containing no genuine tuning, was passed through both reconstruction pipelines and correlated against real model geometry (200 trials): the von Mises pipeline produces mean |*r*| = 0.334 on data with zero real signal, nearly twice the raw pipeline’s |*r*| = 0.168, establishing that the von Mises fitting step independently inflates spurious correlation and should be read as the less trustworthy reference wherever the two disagree.

#### Split-half reliability

Random half-samples of the real Henning cells were independently reconstructed and correlated against each other (500 trials): raw reliability *r* = 0.924 ± 0.026, von Mises *r* = 0.987 ± 0.009, both far above the noise floor above, confirming the dataset carries substantial, genuine, replicable signal regardless of reconstruction method. Both reference versions are reported throughout; where they disagree, raw is treated as the more trustworthy reference, given the fitting artifact established above.

### 2.8 CKA validation

As an independent robustness check on the RSA result (not a replacement for it), we computed linear centered kernel alignment (CKA; Kornblith et al. 2019) between mean real and mean null population matrices, operating on raw activation matrices rather than RDMs:

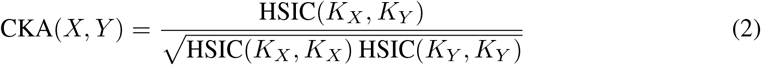

Significance was assessed by stimulus-label permutation test (10,000 permutations); model-level bootstrap 95% confidence intervals were computed by resampling models with replacement (10,000 samples).

### 2.9 Untrained wiring (checkpoint 0)

To test whether real and null wiring are distinguishable prior to any training, each null-scheme network (degree-preserving swap and Erdős–Rényi) was evaluated at checkpoint 0, before any gradient-based optimization, using the identical individual-pairwise RSA methodology (§2.6) applied throughout. This differs from an earlier attempt at this comparison, evaluated at the raw Flyvis prior initialization rather than checkpoint 0 of an actual training run, which failed on a numerical-precision guard rather than yielding an interpretable result; we do not report that earlier construction here, since checkpoint 0 of the null-scheme training run resolves the issue directly and yields a statistically well-resolved comparison.

### 2.10 Trained-random wiring: two independent methods

Both null schemes were trained through the identical recipe described above (§2), *n* = 10 independent networks per scheme, then compared against real, trained wiring two independent ways.

#### Method A: biology-mediated

Each trained network’s RDM was correlated against the Henning biological reference (§2.7), using per-network Mann–Whitney comparison against the real ensemble. Given the real ensemble (*n* = 50) and each null ensemble (*n* = 10) are not naturally size-matched, we additionally resampled the real ensemble down to *n* = 10 (10,000 draws) and reran the comparison on each draw, reporting the median *p*-value and the fraction of draws reaching significance.

#### Method B: direct RDM comparison, no biological reference

Real and null RDMs were compared directly to each other, using the same individual-pairwise methodology as the untrained comparison (§2.9), now at the network’s final, fully-trained checkpoint. This method requires no biological reference at all, and so is immune to any circularity confound in the Henning reference. To characterize how this comparison evolves over the course of training, not just at its two endpoints, we additionally evaluated every checkpoint of every network across the full training run, for all three stimulus conditions.

### 2.11 Training-vs-wiring test

A negative-trending pattern observed in Method A (real wiring correlating *more weakly* with the biological reference than trained null wiring, in one specific comparison) raised a follow-up question: is this about wiring identity, or a property of training itself, independent of which wiring is trained? Two distinct quantities are tested here, and it is worth being explicit that they are not the same measurement. The first is a *snapshot* test: for each model in a population, whether its own single, already-computed correlation against the biological reference (§3.3, Method A) is itself negative rather than positive, i.e., anti-correlated with the reference rather than correlated with it. As throughout this paper, this is a partial correlation with circular structure already removed (§3.2), not a raw correlation, so this test is not confounded by the circularity issue established earlier. Every population tested (real, both trained null schemes, and the original untrained weight-shuffled baseline) was sorted into trained or untrained and compared on this snapshot measure, with the untrained population resampled to match the pooled trained sample size (10,000 draws) given the natural size mismatch. The second is a *trend* test, answering a different question: within a single network’s own training run, does its correlation against the reference systematically rise or fall as training progresses? This is computed by correlating checkpoint index against that network’s own per-checkpoint reference correlation, evaluated across every checkpoint of all 20 trained null-scheme networks, yielding one trend statistic *ρ* per network. A negative *ρ* means that network’s fidelity to the reference declined over the course of its own training; a positive *ρ* means it improved. The snapshot test answers whether trained populations end up negatively correlated; the trend test answers whether individual networks are moving in a consistent direction while they train.

### 2.12 Retraining-based instability test

To test whether the direction of an individual network’s own fidelity trend is determined by its specific wiring realization or by training-process randomness unrelated to wiring, we adopted the retraining methodology of Frankle, Dziugaite, Roy & Carbin (Frankle et al., 2020): holding a network’s wiring fixed and retraining it from scratch with a different training-noise seed, then checking whether the resulting fidelity trend reproduces in sign and magnitude. “Fidelity trend” here is the same *ρ* statistic defined in §2.11, the correlation between checkpoint index and a network’s own per-checkpoint reference correlation, now computed once per retrained replicate rather than once per original network. Eight seed networks (five degree-preserving swap, three Erdős–Rényi) were each retrained twice from identical wiring. A one-way permutation test (20,000 shuffles) compares between-group variance (across the eight distinct wirings) to within-group variance (across replicates of the same wiring): a large, significant F-statistic is consistent with wiring identity determining trend direction; an F-statistic indistinguishable from the permutation null is consistent with training-process randomness.

### 2.13 Reproducibility

All experiments used seed 42 where applicable, with deterministic algorithms enabled. Full code and data availability details are provided in the Data and Code Availability section.

## 3 Results

### 3.1 Untrained real wiring is distinguishable from random wiring, but only once ensemble-mean averaging is replaced by individual-pairwise comparison

Across two stimulus conditions, the ensemble-mean RDM correlation between real (connectome-constrained) and random wiring was *r* = 0.686 (ON edges only, *n* = 50, *p <* 0.0001) and *r* = 0.846 (ON+OFF edges, *n* = 50, *p <* 0.0001). Taken alone these numbers are ambiguous: both are numerically high and could be misread as evidence of similarity rather than difference. The correct comparison is relative, not absolute, how similar real models are to each other versus to random, which the ensemble-mean statistic cannot answer, since it collapses each 50-model ensemble into a single mean RDM before any comparison is made.

The individual-pairwise comparison, every real model’s RDM correlated against every other real model’s RDM (within-real baseline) and against every random model’s RDM (real-vs-random), each set compared by Mann–Whitney *U* test, shows real networks are significantly more similar to each other than to random wiring on both stimulus sets. The ensemble-mean statistic’s understatement of this gap is not merely quantitative: on ON+OFF, the ensemble-mean correlation (*r* = 0.846) numerically exceeds the individual-pairwise within-real baseline itself (*r* = 0.838), a result that is not possible under an honest comparison, since real networks cannot on average be more similar to random wiring than to each other. We therefore report the individual-pairwise statistic as primary throughout.

**Figure 1.**
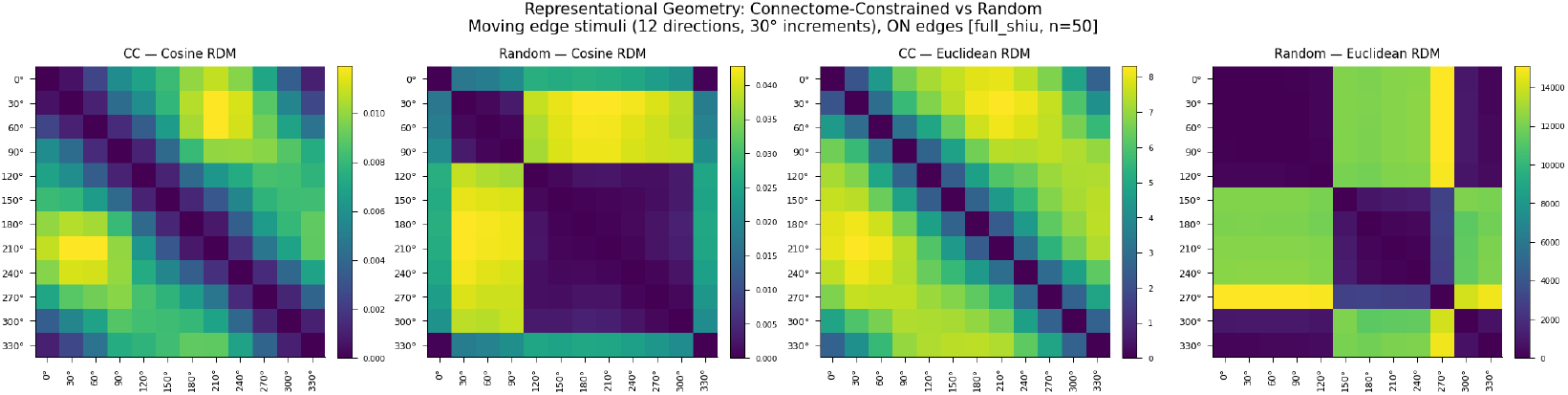
Representational dissimilarity matrices, built from simulated population responses, ON edges only (*n* = 50). Real networks show smooth, graded dissimilarity consistent with continuous direction tuning; random networks show a qualitatively different block structure, splitting responses into two large clusters rather than a graded direction-tuned pattern.

**Figure 2.**
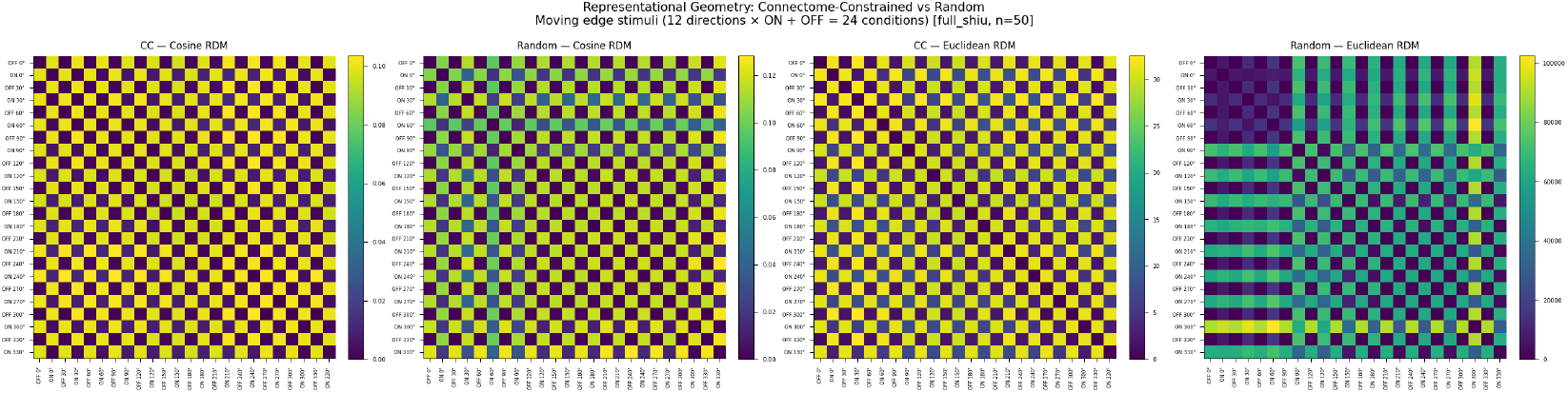
Representational dissimilarity matrices, ON+OFF edges (*n* = 50). Real networks show a sharp, regular checkerboard pattern reflecting consistent ON/OFF discrimination across all directions; random networks show this structure only weakly (cosine) or not at all (Euclidean).

#### This distinction holds for the topology-randomizing null schemes items 3–5 depend on, not just the weight-shuffled baseline

Degree-preserving swap and Erdős–Rényi were each evaluated at checkpoint 0 (before any training), using the identical individual-pairwise methodology, across three stimulus conditions.

Every comparison reaches significance, but the size of the gap varies considerably in a consistent pattern: ON-only and the independently-built Henning 8-direction set agree closely with each other, both showing degree-preserving swap essentially uncorrelated with real wiring and Erdős–Rényi showing a modest but real relationship. ON+OFF stands apart, showing a much smaller gap for both null schemes, still significant, but qualitatively different.

#### This turns out to have a specific, identifiable cause

The ON+OFF RDM contains a component ON-only and Henning structurally cannot have, whether a stimulus is ON- or OFF-polarity, a coarse distinction plausibly reproduced by any network, real or random. Decomposing the ON+OFF RDM into its ON–ON and OFF–OFF sub-blocks and discarding the cross-polarity terms collapses both null schemes’ apparent convergence, landing almost exactly on the independently-run pure ON-only result (Table 3).

**Table 1:** Individual-pairwise comparison recovers a real-versus-random gap the ensemble-mean statistic understates. All values from simulated-response RDMs (cosine distance), weight-shuffled random baseline, *n* = 50 per condition.

| Stimulus set | Ensemble-mean $r$ | Within-real $r$ | Real-vs-random $r$ | Mann–Whitney $p$ |
| --- | --- | --- | --- | --- |
| ON-only | 0.686 | 0.721 | 0.404 | $1.7 \times 10^{-304}$ |
| ON+OFF | 0.846 | 0.838 | 0.754 | $1.2 \times 10^{-141}$ |

**Table 2:** Real wiring is distinguishable from both topology-randomizing null schemes, untrained, across all three stimulus conditions. *n* = 10 models per scheme.

| Stimulus set | Scheme | Within-real $r$ | Real-vs-null $r$ | Mann–Whitney $p$ |
| --- | --- | --- | --- | --- |
| ON-only (12 dir) | Degree-preserving swap | $0.838 \pm 0.078$ | $-0.032 \pm 0.109$ | $3.50 \times 10^{-22}$ |
| ON-only (12 dir) | Erdős–Rényi | $0.838 \pm 0.078$ | $0.264 \pm 0.149$ | $3.50 \times 10^{-22}$ |
| ON+OFF (24 cond.) | Degree-preserving swap | $0.850 \pm 0.057$ | $0.764 \pm 0.040$ | $1.33 \times 10^{-13}$ |
| ON+OFF (24 cond.) | Erdős–Rényi | $0.850 \pm 0.057$ | $0.794 \pm 0.048$ | $8.72 \times 10^{-8}$ |
| Henning 8-direction | Degree-preserving swap | $0.792 \pm 0.101$ | $-0.089 \pm 0.139$ | $3.49 \times 10^{-22}$ |
| Henning 8-direction | Erdős–Rényi | $0.792 \pm 0.101$ | $0.186 \pm 0.149$ | $3.80 \times 10^{-22}$ |

**Table 3:** Within-polarity decomposition recovers the ON-only result, confirming the full ON+OFF comparison’s smaller gap was a confound. Untrained.

| Scheme | Comparison | Real-vs-null $r$ | Mann–Whitney $p$ |
| --- | --- | --- | --- |
| Degree-preserving swap | Full ON+OFF | $0.762 \pm 0.039$ | $5.09 \times 10^{-14}$ |
| Degree-preserving swap | ON–ON sub-block | $-0.040 \pm 0.117$ | $3.49 \times 10^{-22}$ |
| Degree-preserving swap | OFF–OFF sub-block | $-0.010 \pm 0.135$ | $4.13 \times 10^{-22}$ |
| Erdős–Rényi | Full ON+OFF | $0.793 \pm 0.048$ | $9.79 \times 10^{-8}$ |
| Erdős–Rényi | ON–ON sub-block | $0.259 \pm 0.149$ | $3.50 \times 10^{-22}$ |
| Erdős–Rényi | OFF–OFF sub-block | $0.319 \pm 0.190$ | $3.93 \times 10^{-14}$ |

**Table 4:** The Henning reference is substantially less circular than the retired Maisak construction, though not negligibly circular itself.

| Reference | Variance explained by circular structure ($r^2$) |
| --- | --- |
| Henning, raw | 63.5% |
| Henning, von Mises | 82.3% |
| Maisak | 95.6% |

The apparent convergence in the full ON+OFF comparison was real, but it was measuring shared ON/OFF-discrimination ability, not a genuine loss of fine-grained direction-tuning distinctiveness. ON+OFF is not a different result from ON-only and Henning; it is the same result diluted by a confound. One further asymmetry surfaced by this decomposition, not yet explained: real wiring’s own within-ensemble consistency is itself lower on OFF-only structure (*r* = 0.634) than ON-only structure (*r* = 0.838), independent of any null-scheme comparison.

**Figure 3.**
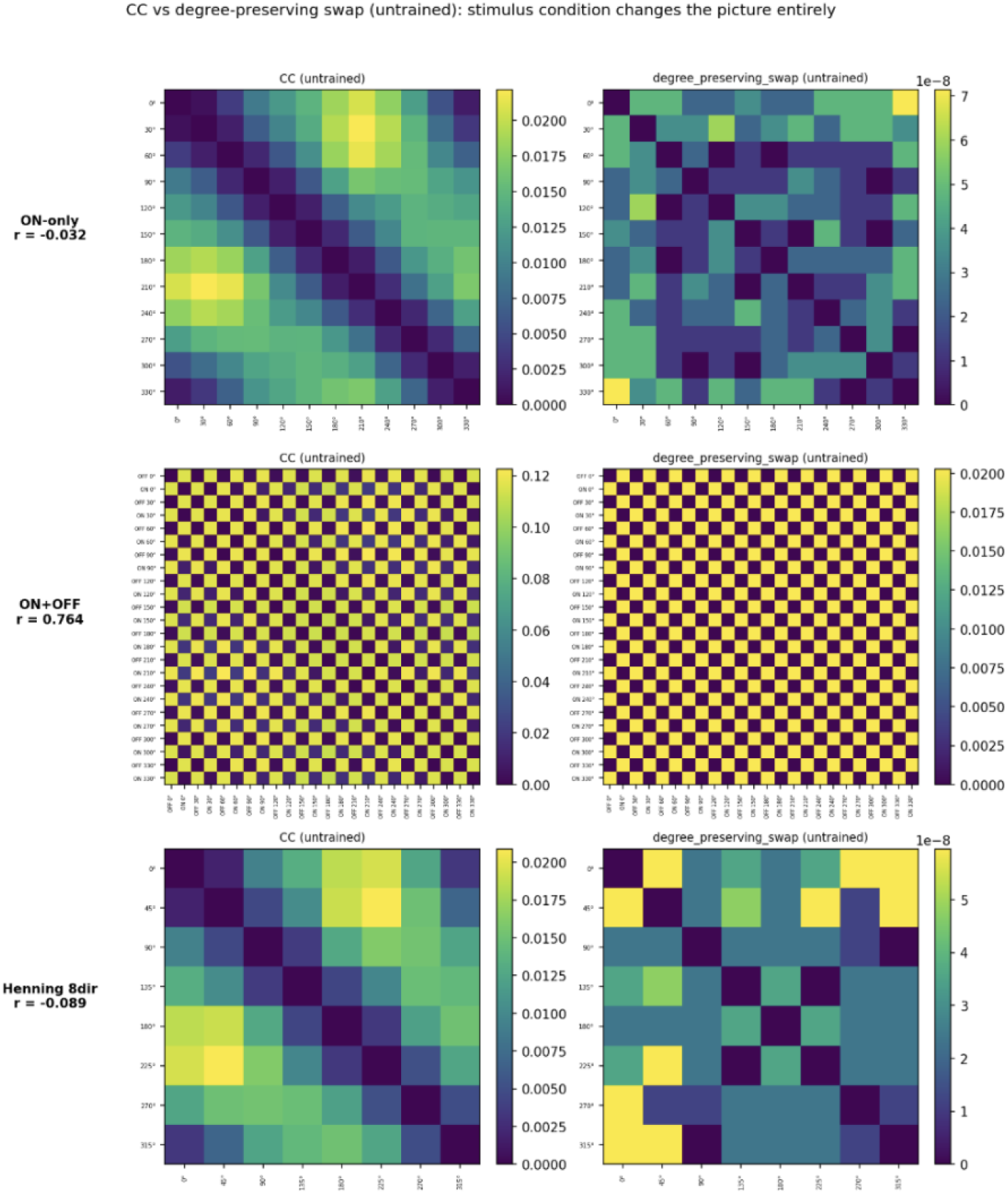
Real (left) vs. untrained degree-preserving-swap wiring (right), three independently-built stimulus conditions. ON-only (top, *r* = −0.032) and Henning 8-direction (bottom, *r* = −0.089) each show real and null wiring as essentially unrelated. ON+OFF (middle, *r* = 0.764) shows a much smaller gap, real and null wiring look considerably more alike once OFF-polarity stimuli are included, because both reproduce the coarse ON/OFF-discrimination signal. Since the two independently-built stimulus sets (ON-only, Henning) agree closely, ON+OFF is the condition that stands apart, not the other way around.

#### Answer

Yes: geometry distinguishes real from random wiring when untrained, for both topology-randomizing null schemes Experiments 3–5 depend on, not just the original weight-shuffled baseline. ON+OFF’s smaller gap is not a real exception; it reflects dilution by shared ON/OFF-discrimination structure and disappears once within-polarity structure is isolated.

### 3.2 The original biological reference is confounded with circular stimulus structure; a validated replacement is substantially, but not entirely, non-circular

The stimulus set throughout this work is a set of directions arranged around a compass. Purely from that layout, without any biology involved, nearby directions look more similar than distant ones, a predictable, purely geometric pattern we refer to as circular structure. Because almost any model, biologically accurate or not, will trivially reproduce this pattern (it is a property of the stimulus layout, not of any specific computation), a biological reference dominated by circular structure cannot discriminate a faithful model from an unfaithful one: both would score well, for the wrong reason.

The original reference, built from idealized von Mises curves fit to summary parameters in Maisak et al. (Maisak et al., 2013), is dominated by exactly this confound. Maisak’s own data show T5 cells largely fail to respond to moving ON-edge stimuli; on the ON-only stimulus set, this collapses the eight nominal T4/T5 subtypes to four T4 curves of identical width at even 90^°^ spacing, a shape whose pairwise distances are almost entirely explained by angular position alone (*r* = 0.978 against a pure circular-distance model). After partialling out circular structure, the residual signal was no longer significant at *n* = 50. This invalidated the original biology-mediated comparison and the first version of the trained-random test that depended on the same reference.

We instead constructed the biological reference from Henning, Ramos-Traslosheros, Gür & Silies (Henning et al., 2022), real per-cell T4/T5 recordings rather than idealized summary curves, in two independent ways: a raw version (each cell’s peak response averaged directly, no curve-fitting) and a von Mises version (each cell’s own fitted preferred direction and concentration, unlike the retired construction’s single shared curve per layer).

Two further tests establish that the non-circular remainder is genuine signal, not reconstruction artifact. First, pure synthetic noise, containing no genuine tuning, was passed through both reconstruction pipelines and correlated against real model geometry (Table 5): the von Mises pipeline produces spurious correlation nearly twice the magnitude of the raw pipeline’s, on data with zero real signal, establishing that von Mises independently inflates correlation and should be treated as the less trustworthy reference wherever the two disagree.

**Table 5:**
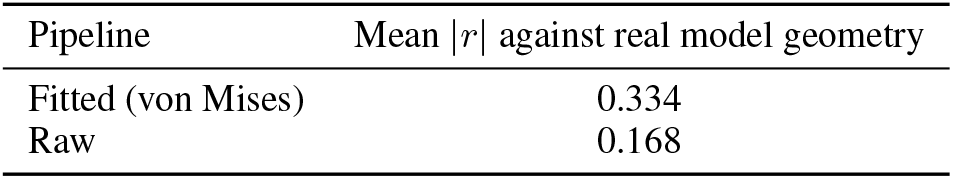
Curve-fitting inflates spurious correlation on pure noise. 200 trials.

Second, random half-samples of the real Henning cells were independently reconstructed and correlated against each other (Table 6): both versions show reliability far above the noise floor above, confirming the dataset carries substantial, genuine, replicable signal regardless of reconstruction method.

**Table 6:** Split-half reliability confirms genuine signal in the real Henning data. 500 trials.

| Pipeline | Reliability ($r$) | Std. across trials |
| --- | --- | --- |
| Fitted (von Mises) | 0.987 | 0.009 |
| Raw | 0.924 | 0.026 |

**Table 7:**
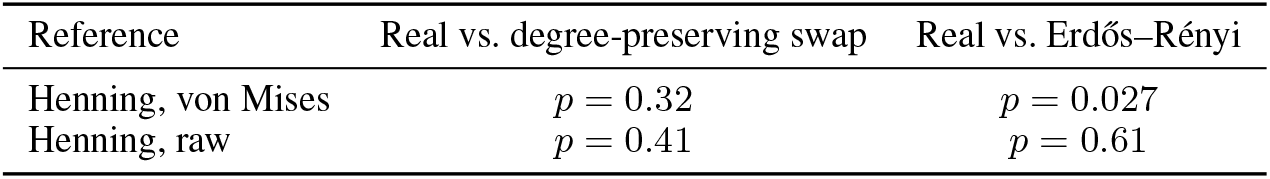
Real vs. trained-random wiring, unmatched sample sizes.

| Reference | Real vs. degree-preserving swap | Real vs. Erdős–Rényi |
| --- | --- | --- |
| Henning, von Mises | $p = 0.32$ | $p = 0.027$ |
| Henning, raw | $p = 0.41$ | $p = 0.61$ |

**Table 8:** Size-matched comparison: the degree-preserving null is a genuine null; the Erdős– Rényi result does not survive. Median *p* across 10,000 draws; % of draws reaching *p <* 0.05 in parentheses.

| Reference | Real vs. degree-preserving swap | Real vs. Erdős–Rényi |
| --- | --- | --- |
| Henning, von Mises | $p = 0.47$ (0%) | $p = 0.09$ (30%) |
| Henning, raw | $p = 0.52$ (0.4%) | $p = 0.65$ (0.2%) |

**Table 9:** Within-polarity decomposition, trained. Unlike untrained (Table 3), isolating within-polarity structure does not fully restore separation for both schemes.

| Scheme | Comparison | Real-vs-null $r$ | Mann–Whitney $p$ |
| --- | --- | --- | --- |
| Degree-preserving swap | ON–ON sub-block | $0.695 \pm 0.130$ | $5.87 \times 10^{-10}$ |
| Degree-preserving swap | OFF–OFF sub-block | $0.494 \pm 0.175$ | $1.22 \times 10^{-5}$ |
| Erdős–Rényi | ON–ON sub-block | $0.707 \pm 0.123$ | $1.92 \times 10^{-9}$ |
| Erdős–Rényi | OFF–OFF sub-block | $0.592 \pm 0.165$ | 0.087 |

#### Answer

No, not entirely: the original reference measured circular stimulus structure more than direction tuning. The Henning dataset is still substantially circular, but has a meaningfully smaller circular component than Maisak, and the non-circular remainder is independently validated, via split-half reliability tested against a synthetic-noise floor, as genuine, reproducible signal rather than reconstruction artifact. Both versions are reported throughout what follows; where they disagree, raw is treated as the more trustworthy reference.

### 3.3 Trained real wiring is largely indistinguishable from trained random wiring, by two independent methods

With a validated non-circular reference in hand, we tested the specific limitation Brunton et al. (Brunton et al., 2026) name but do not test themselves: they predict training the connectome’s synaptic weights would only deepen the interpretability problem their untrained-random result raises. Does representational geometry distinguish real, trained wiring from random wiring that has also been trained to task adequacy?

#### Method A: biology-mediated

Each trained null scheme (*n* = 10) was correlated against the Henning reference and compared to real, trained wiring (*n* = 50) by per-network Mann–Whitney test.

Against the degree-preserving null, real and random wiring are statistically indistinguishable on both references. Against Erdős–Rényi, the same holds on raw, but not on von Mises, where the difference is significant and runs in the direction opposite a fidelity hypothesis: random wiring shows a *stronger* relationship with the reference than real wiring does. Given the real (*n* = 50) and null (*n* = 10) ensembles are not naturally size-matched, we resampled the real ensemble down to *n* = 10 (10,000 draws) and reran the comparison on each draw.

The degree-preserving result holds up, genuinely null, not just null by small sample. The Erdős–Rényi result does not survive the fair comparison: median *p* rises from 0.027 to 0.09, and only 30% of size-matched draws reach significance at all. The original result was, at least in part, an artifact of the sample-size asymmetry; the fact that it appeared only on the von Mises reference (never raw), combined with von Mises’s independently confirmed inflation of spurious correlation (§3.2), provides a second, independent reason for caution.

#### Method B: direct RDM comparison, no biological reference

Real and null RDMs were compared directly, using the individual-pairwise methodology of §3.1, at each network’s final checkpoint, and at every checkpoint across the full training run, for all three stimulus conditions.

**Figure 4.**
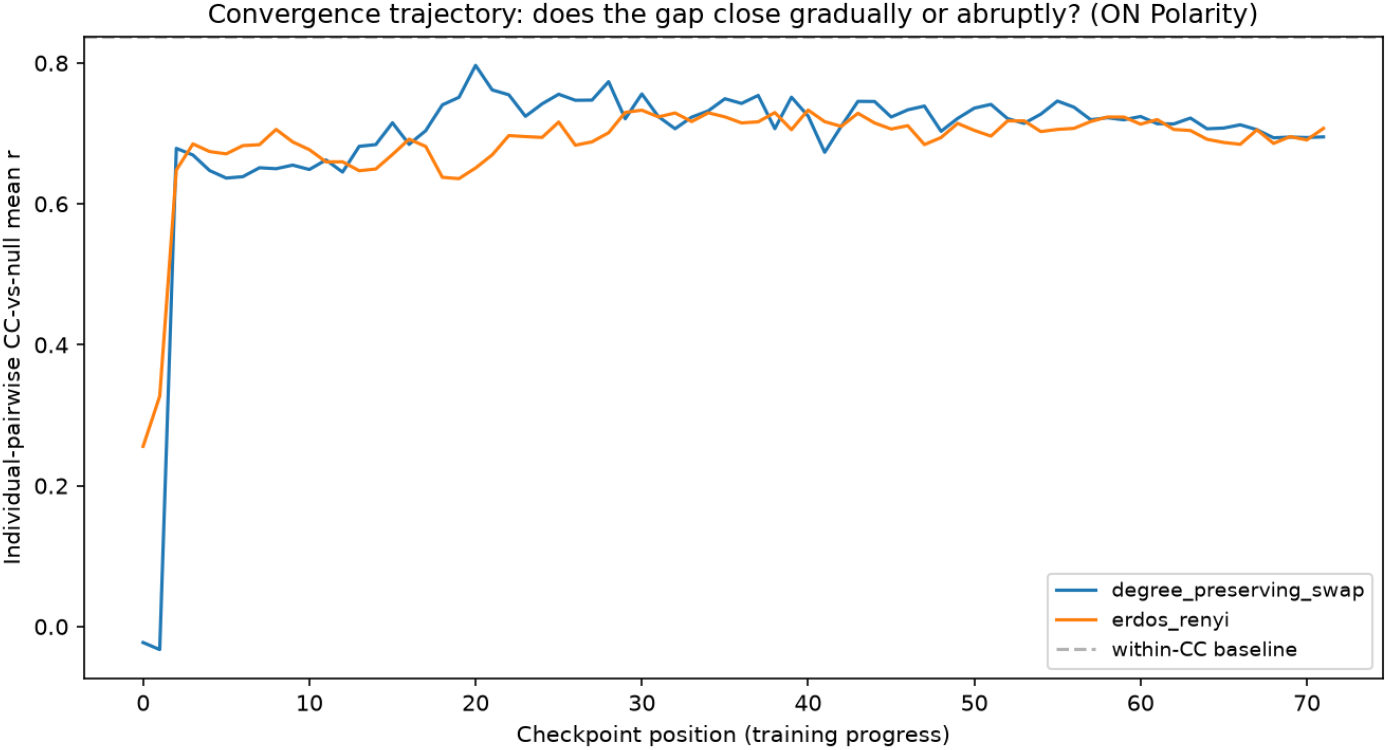
Convergence trajectory, ON-only. Starting from essentially no relationship (checkpoint 0: − 0.023 degree-preserving, +0.256 Erdős–Rényi), both schemes rise sharply within the first 10–20 checkpoints, then plateau in a noisy band (final: 0.695 and 0.707, against a within-real baseline of 0.838).

**Figure 5.**
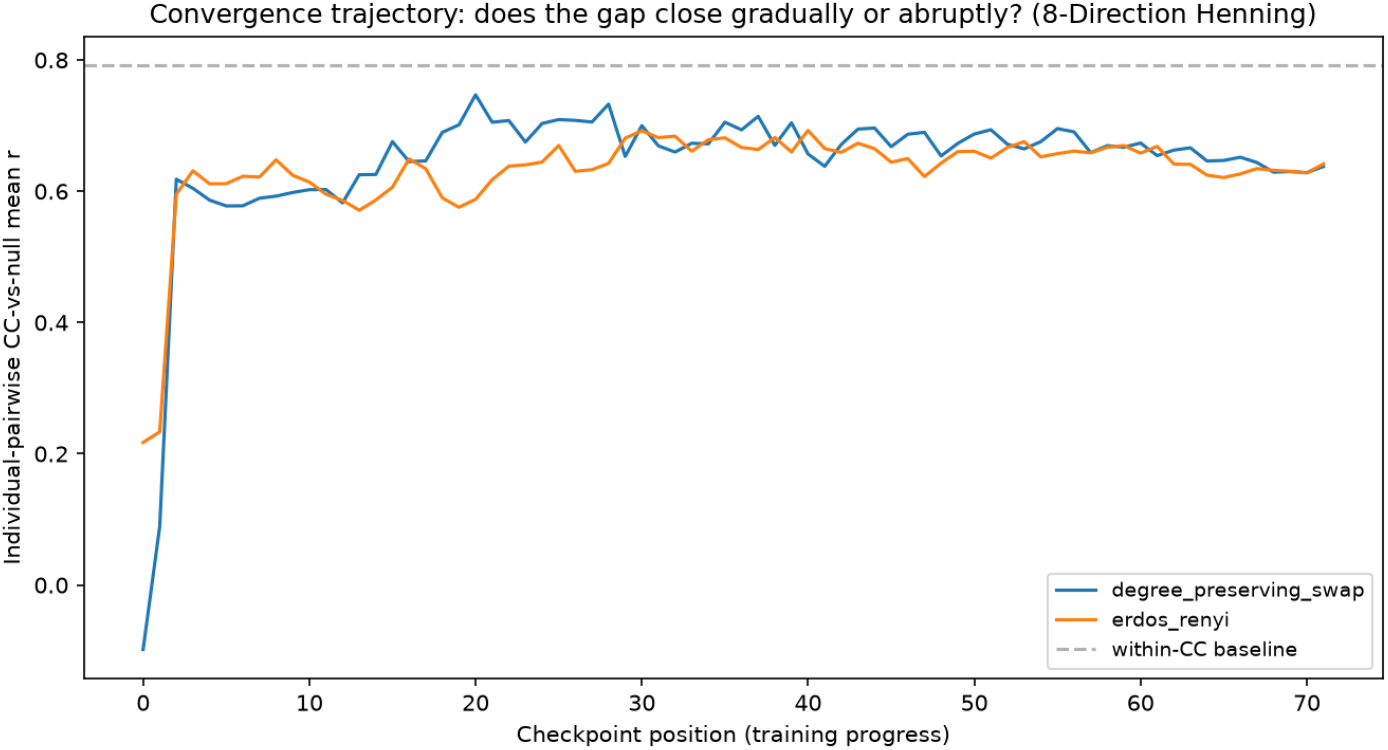
Convergence trajectory, Henning 8-direction. The same pattern as ON-only: near-zero start, abrupt rise within the first 2–3 checkpoints, then a long, stable plateau (0.58–0.75) for the remaining ∼86% of training, well short of the baseline (∼0.79).

**Figure 6.**
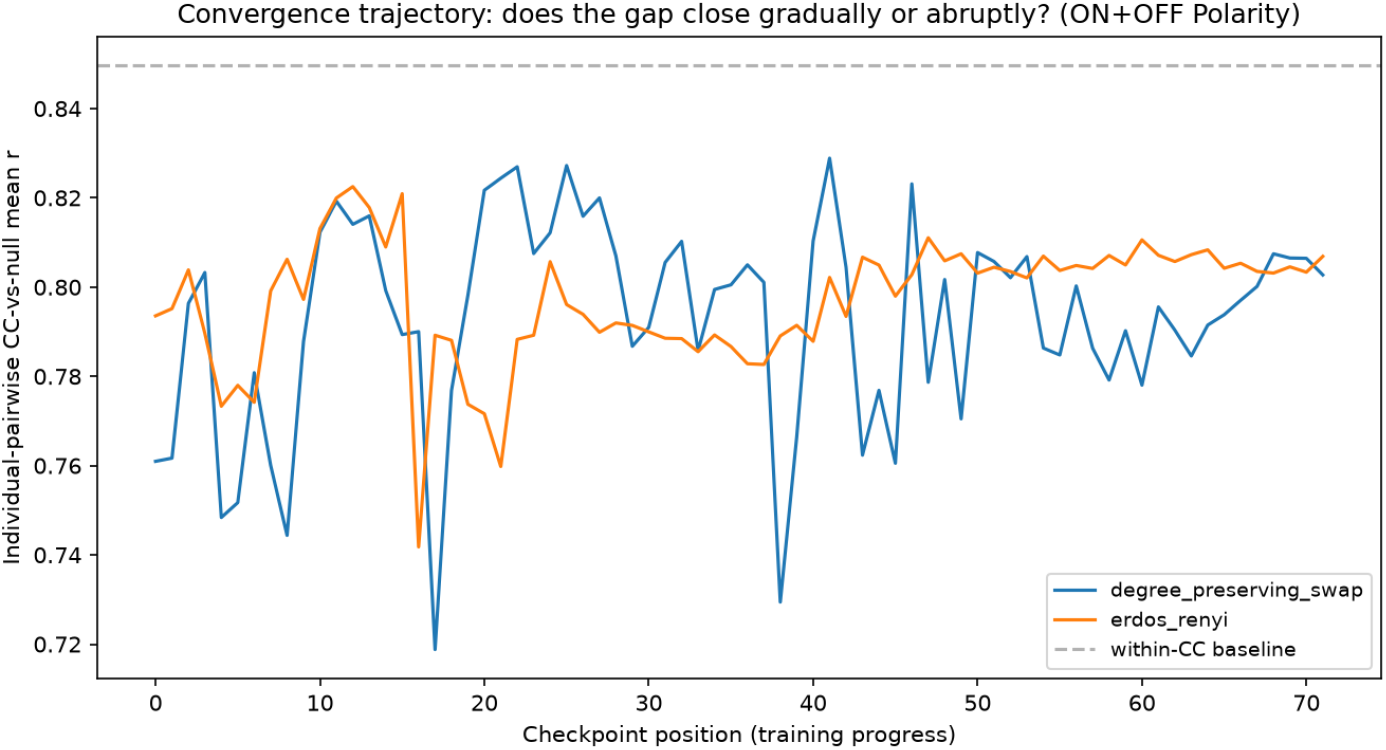
Convergence trajectory, ON+OFF. No jump: both schemes start already near their final level (checkpoint 0: 0.761 degree-preserving, 0.794 Erdős–Rényi) and simply oscillate in that range for the entire run (final: 0.803 and 0.807, baseline 0.850).

All three trajectories resolve to one unified rule: the shape of a network’s convergence trajectory is set entirely by how much distinguishability existed at checkpoint 0, not by wiring scheme, not by a separate gradual training process. ON-only and Henning both started with essentially no untrained relationship and both show the same abrupt-jump-then-plateau shape; ON+OFF started already close to its final value and shows no jump at all. Neither gradual convergence nor a non-monotonic process describes any of the three conditions; it is fast equilibration followed by stable noise, and how large the initial jump looks depends entirely on the starting distance.

The ON–ON sub-block, extracted from the pooled trained data, matches the standalone trained ON-only result almost exactly (degree-preserving: 0.695 vs. 0.695; Erdős–Rényi: 0.707 vs. 0.706), confirming the decomposition behaves consistently. Trained OFF–OFF, however, tells a different story than untrained: it stays significant for degree-preserving swap but loses significance entirely for Erdős–Rényi, the first non-significant comparison anywhere in this analysis. The untrained confound explanation (§3.1) does not contradict this, it is specific to untrained wiring; once training happens, real convergence occurs within each sub-block independently, and for Erdős–Rényi on OFF-polarity structure specifically, that convergence reaches full statistical indistinguishability. Checking every checkpoint of training confirms this is not an early or gradual effect: the OFF–OFF comparison stays significant for nearly the entire trajectory and only crosses into non-significance at the final checkpoint, leaving open whether this reflects a genuine, late-emerging effect or a boundary case.

#### Answer

No, largely, and two independent methods agree. Method A: once properly size-matched, real and trained-random wiring are statistically indistinguishable from both null schemes, on both references. Method B: training substantially narrows the gap wherever an untrained gap existed, and in one specific case, trained Erdős–Rényi on OFF-polarity structure, closes it completely, the only comparison across either method where that happens outright. The broader convergence happens almost immediately at the start of training, not gradually across the run.

### 3.4 The negative-trending pattern is driven by training, not by wiring identity

Method A’s Erdős–Rényi result, before its resampling correction, pointed in the wrong direction for a fidelity story: random wiring correlated more strongly with the biological reference than real wiring did. We tested whether this reflects wiring identity or a property of training itself, independent of which wiring is trained.

The first test is a snapshot: for each model in a population, is its own single correlation against the biological reference (already computed for Method A) itself negative, meaning that model’s representational geometry is anti-correlated with the reference rather than correlated with it. “% negative” in Table 10 is the fraction of models in that population for which this is true.

**Table 10:** Trained populations trend negative against the von Mises reference; the untrained population does not. “% negative” is the fraction of models in each population whose own single correlation against the reference is itself negative. Degree-preserving swap and Erdős–Rényi reflect updated evaluations at *n* = 25 and *n* = 24 respectively; one Erdős–Rényi network was excluded as unstable (a constant, non-direction-selective output, all pairwise RDM distances exactly zero), following the same exclude-and-report stability discipline used throughout this paper. The pooled row is recomputed at these updated sample sizes.

| Population | Trained? | $n$ | % negative | Significance |
| --- | --- | --- | --- | --- |
| Real | Yes | 50 | 78% | $p = 0.0001$ |
| Erdős–Rényi | Yes | 24 | 67% | n.s. ($p = 0.152$) |
| Degree-preserving swap | Yes | 25 | 76% | $p = 0.0146$ |
| Pooled trained (both schemes) | Yes | 49 | 71% | $p = 0.0038$ |
| Weight-shuffled random | No | 50 | 54% | n.s. (chance) |

**Table 11:** Permutation test, replicate *ρ* as within-vs. between-group scatter. A large, significant *F* is consistent with wiring identity determining trend direction.

| Scope | von Mises $F$ | von Mises $p$ | Raw $F$ | Raw $p$ |
| --- | --- | --- | --- | --- |
| Degree-preserving swap ($n = 5$) | 4.88 | 0.025 | 1.34 | 0.311 |
| Erdős-Rényi ($n = 3$) | 1.53 | 0.218 | 0.93 | 0.492 |
| Pooled ($n = 8$) | 4.93 | 0.006 | 1.16 | 0.370 |

The cleanest single comparison, real vs. weight-shuffled random, matched in size (*n* = 50 each), is significant on the raw reference (Mann–Whitney *p* = 0.008). Given the pooled trained comparison’s own size mismatch (*n* = 20 vs. *n* = 50), the untrained population was resampled to *n* = 20 (10,000 draws): median *p* = 0.0077 (IQR 0.0028–0.0193), with 92.7% of draws reaching significance, confirming the finding does not rest on the size mismatch. One further check, applied to every population in Table 10, is worth reporting directly: whether the ensemble-mean correlation (averaging RDMs before correlating) and the per-network mean (averaging already-computed per-network correlations) actually agree, the same ensemble-mean-versus-individual-pairwise concern raised in §3.1, applied here to this different statistic. For weight-shuffled random specifically, they diverge on both references, with the ensemble-mean and per-network mean disagreeing in sign on both, not just in magnitude. Its snapshot result in Table 10 is reported as is, but should be read with that caveat: the ensemble-level summary for this population is not representative of how its individual networks actually behave.

**Figure 7.**
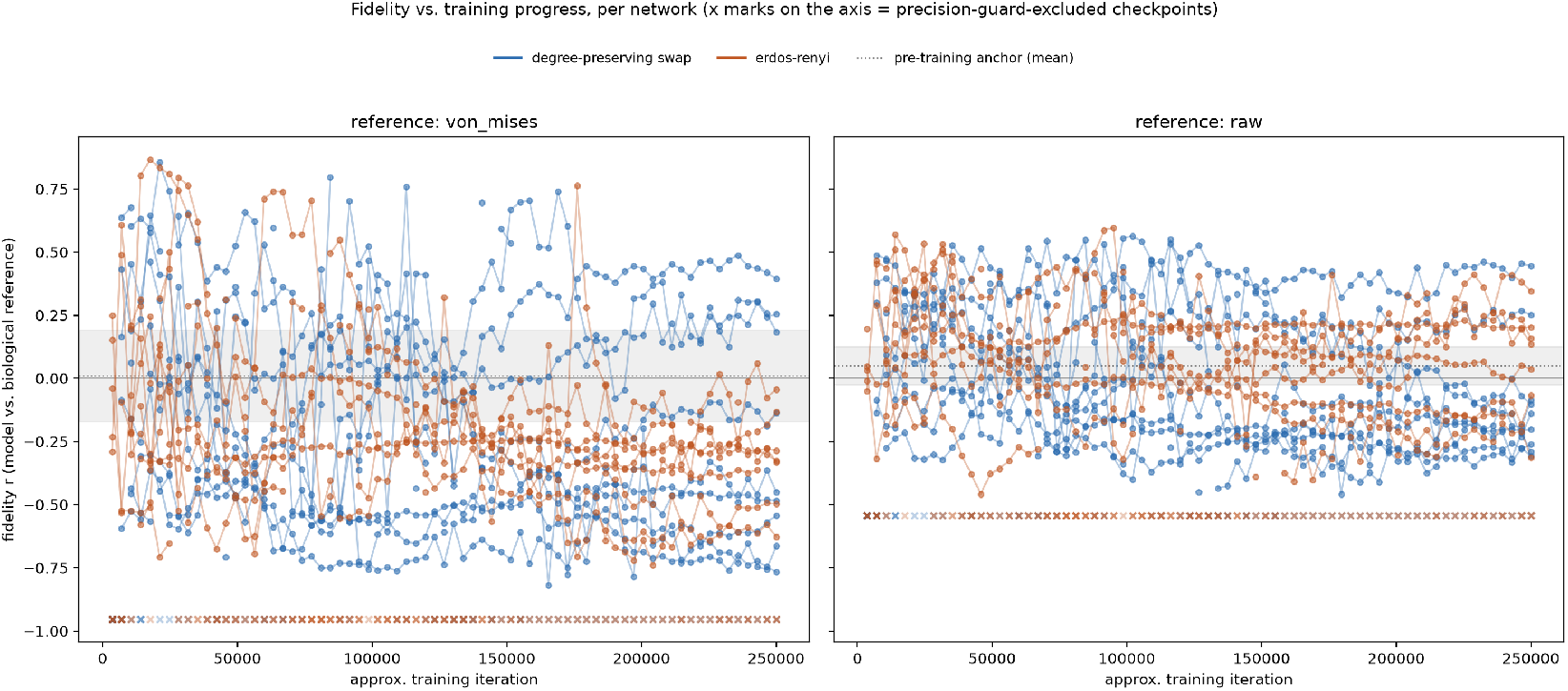
Fidelity vs. training progress, all 20 trained null-scheme networks, both references. X marks indicate checkpoints excluded by a precision guard. Individual trajectories are visibly coherent, not random jitter, but fan out in both directions with no shared population-level trend.

The second test asks a different question, not whether a model’s endpoint correlation is negative, but whether that correlation moves in a consistent direction as a given network trains. For each of the 20 trained null-scheme networks, we correlate checkpoint index against that network’s own per-checkpoint correlation to the biological reference, yielding one trend statistic *ρ* per network: *ρ <* 0 means that network’s fidelity to the reference declined over its own training; *ρ >* 0 means it improved. Evaluating all 20 trained networks at every checkpoint of their own training reveals a real effect within nearly every individual network, several reach |*ρ*| *>* 0.7–0.9 at *p <* 0.0001, but the direction is idiosyncratic: within the degree-preserving scheme alone, four networks trend strongly negative while five trend strongly positive. A sign test across all 20 finds no consistent population-level direction (11*/*20 negative, *p* = 0.82 von Mises; 13*/*20 negative, *p* = 0.26 raw), while a combined significance test is overwhelming on both references (*χ*^2^ = 260.4, *p <* 0.0001 von Mises; *χ*^2^ = 250.3, *p <* 0.0001 raw), evidence something real happens within nearly every network, just not the same something. This is not in tension with the snapshot test above; it is a different measurement of a different question, a majority-negative endpoint (Table 10) can coexist with individual within-network trends diverging in direction along the way.

#### Answer

(a) Yes, this is about training, not wiring; training makes a network more likely to trend negative, and this holds up cleanly across a fair, size-matched comparison. (b) Yes, individual networks shift strongly over their own training, but in different directions from each other, ruling out a simple shared trend and raising the question addressed next: what determines which direction a given network goes?

### 3.5 No significant evidence that wiring identity, rather than training-process randomness, determines an individual network’s fidelity-trend direction

To test whether an individual network’s fidelity-trend direction is determined by its specific wiring realization or by training-process randomness, we adopted the retraining methodology of Frankle, Dziugaite, Roy & Carbin (Frankle et al., 2020): holding wiring fixed and retraining from scratch with a different training-noise seed, then checking whether the resulting trend reproduces in sign and magnitude. Eight seed networks (five degree-preserving swap, three Erdős–Rényi) were each retrained twice.

**Figure 8.**
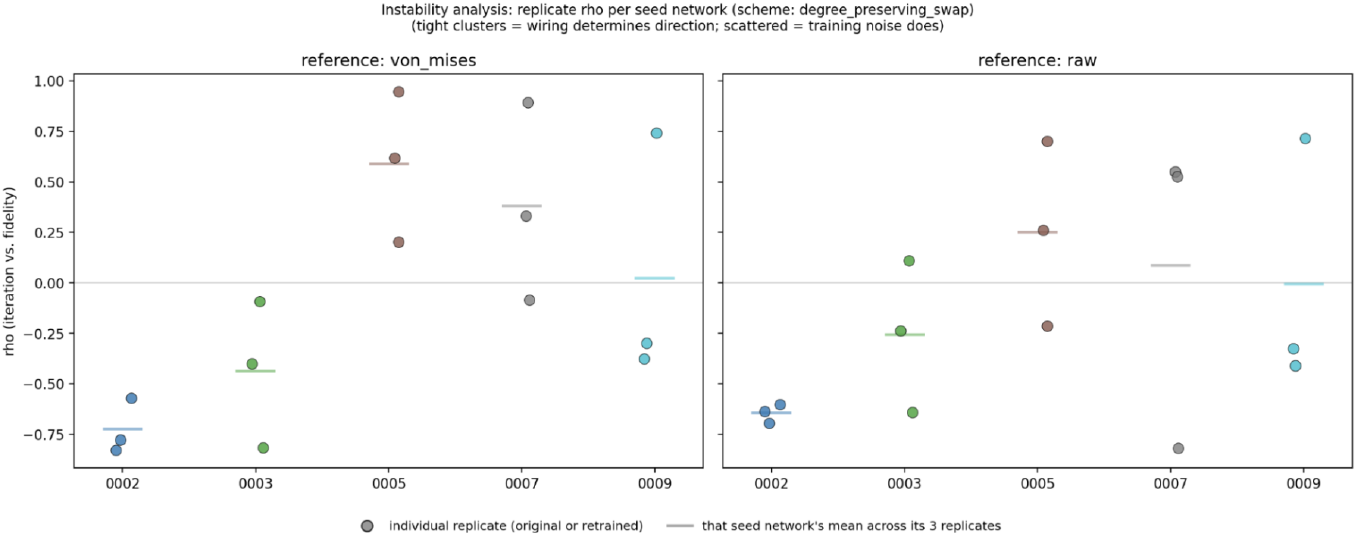
Replicate *ρ* per seed network, both references, degree-preserving swap. 0002 clusters tightly; 0003 and 0005 show internal scatter but stay same-signed as their own original.

**Figure 9.**
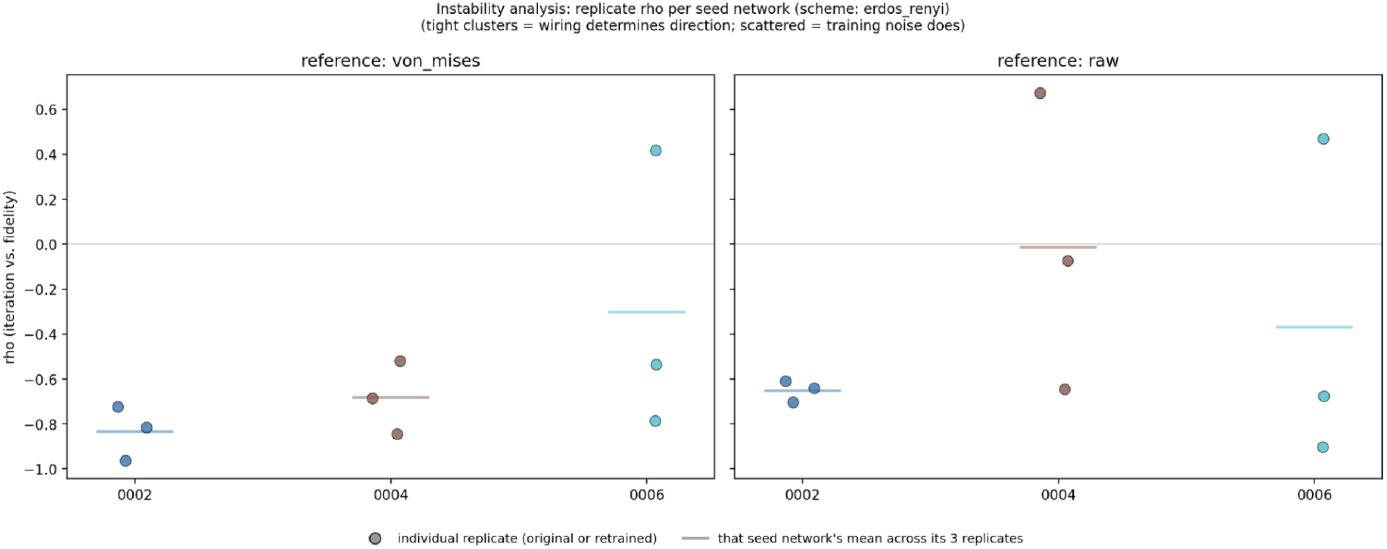
Replicate *ρ* per seed network, both references, Erdős–Rényi. 0002 shows the tightest clustering of any network in this experiment, especially on raw; 0004 and 0006 scatter, with the two references disagreeing sharply within the same network.

Two networks show internal dynamics beyond simple clustering or scatter. 0007’s own original trajectory reverses direction mid-training (first-half *ρ* = −0.819, second-half +0.736); its near-zero full-trajectory value was never a real flat signal but two opposing trends canceling. Its replicates diverge further: seed1 also reverses, in the mirror-image temporal order, while seed2 shows no reversal at all, three qualitatively different dynamics from one wiring. 0009 shows no such internal instability in its own trajectory (consistently positive, +0.618 then +0.257), yet both retrained replicates independently show the same distinctive shape, an early swing opposite the original’s direction followed by partial recovery. Neither pattern is explained by the aggregate test below; 0007 in particular resembles a wiring sitting near a genuine bifurcation.

Von Mises reaches significance twice; raw, already established as the more trustworthy reference (§3.2), does not reach significance in any of the three tests. Every von Mises significant result fails to replicate on raw, consistently, at every level of aggregation.

#### Answer

On the more trustworthy reference, no significant evidence that wiring realization determines the direction of a network’s fidelity trend, in either scheme, alone or combined. This leans toward training-process randomness, though genuine heterogeneity, some networks cluster tightly, others scatter or flip entirely, means the test may simply remain underpowered at *n* = 8 rather than cleanly resolving in either direction.

### 3.6 CKA corroborates the untrained RSA result

As an independent robustness check on §3.1, not a replacement for it, linear CKA (Kornblith et al., 2019) between mean real and mean random population matrices gave Experiment 1 (ON edges) CKA = 0.502 (*p* = 0.0095, bootstrap 95% CI [0.412, 0.781]) and Experiment 2 (ON+OFF edges) CKA = 0.647 (*p <* 0.0001, bootstrap 95% CI [0.052, 0.753]), both significantly greater than chance by stimulus-label permutation test.

**Figure 10.**
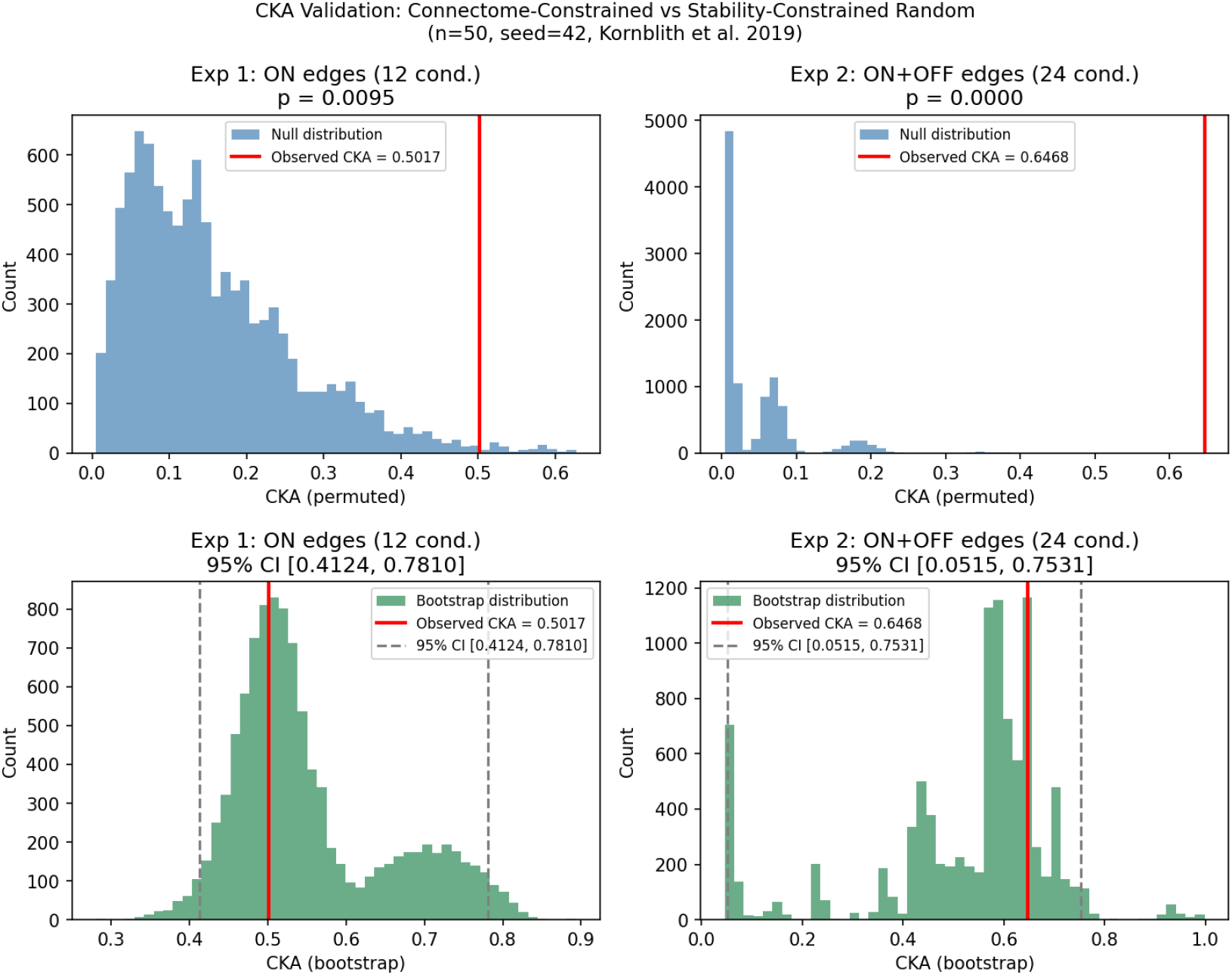
CKA corroborates the RSA fidelity signal. *Top:* permutation null distributions. *Bottom:* model-level bootstrap distributions. Observed CKA (red line) falls outside the permutation null in both conditions.

The convergence of two independent similarity metrics, RSA operating on RDMs and CKA operating on raw activation matrices, strengthens the untrained-distinguishability result of §3.1. The Experiment 2 bootstrap CI is wide due to a bimodal distribution from near-overflow activations in some resampled random models; the permutation test is the primary inference here.

## 4 Discussion

We have shown that representational geometry discriminates real from random wiring across three random-wiring families and every stimulus condition tested, before training, once the correct statistic is used: ensemble-mean comparison understates this gap and in one condition inverts it entirely, while individual-pairwise comparison, corroborated independently by CKA, resolves it cleanly. Grounding this claim in measured biology required repairing the reference itself, the original T4/T5 reference is 97.8% explained by circular stimulus structure and cannot distinguish direction-tuning fidelity from circular ordering; a validated replacement built from real per-cell recordings is substantially, though not entirely, non-circular, and carries genuine signal by independent tests against a noise floor and via split-half reliability.

### The trained-wiring question, largely answered

The core finding extends Brunton et al. (Brunton et al., 2026) in a direction their own paper names but does not test. They demonstrate that a worm connectome trained to control a fly body is behaviorally adequate yet biologically arbitrary, and explicitly predict, without testing, that training the connectome’s own synaptic weights would only deepen this interpretability problem. We tested that prediction directly, and the honest answer is largely, but not entirely, confirmatory of it. A biology-mediated comparison finds real and trained-random wiring statistically indistinguishable, once fairly size-matched, for both null schemes tested. A second, independent method requiring no biological reference at all finds the same substantial narrowing wherever an untrained gap existed, and in one specific case, trained Erdős–Rényi wiring on OFF-polarity structure, finds the gap closes completely. Two independent methods converging on largely the same answer through different routes is real evidence that training, not wiring identity, is what drives this loss of distinguishability once both sides are compared fairly. This is not the confirmation we set out to find, that biological wiring’s geometric signature survives training and therefore constitutes a fidelity metric behavioral benchmarks cannot provide, but we regard the honest null, reached by a method built specifically to test it, as the more useful result. Checkpoint-level trajectories add a further, unanticipated finding: wherever convergence occurs, it happens almost immediately at the start of training and then holds steady, not as a gradual erosion across the full run, so the size of the untrained gap, not the duration of training, is what determines how much distinguishability survives. Concretely, this window is narrow relative to the full training run: the sharp rise completes within the first 10–20 of 72 evaluated checkpoints for ON-only, and within just 2–3 checkpoints for Henning, after which the trajectory holds a stable plateau for roughly the remaining 86% of training. This pattern is consistent with the broader critical-learning-periods literature in deep networks (Achille et al., 2019), which similarly finds that a brief early transient, small relative to total training length, determines much of a network’s eventual solution, after which further training moves the network comparatively little; our checkpoint-level trajectories may be a domain-specific instance of that same general phenomenon rather than something unique to this task. To be explicit about the scope of this connection: we claim only that the observed trajectory shape matches this broader pattern, not that the same underlying mechanism is responsible. Achille et al. attribute their critical periods to the Fisher Information of the network’s weights rising and then falling as the network crosses a curvature bottleneck early in training; we have not measured Fisher Information or curvature in this system, and make no claim that the same mechanism explains our result. A contemporaneous connectome-simulation study of the fly ventral nerve cord offers a useful point of contrast. Pugliese et al. (Pugliese et al., 2025) found that structural connectivity alone, using randomly drawn, unfit biophysical parameters rather than the task-optimization procedure used throughout this paper, was sufficient to identify a minimal central pattern generator circuit for walking, a prediction they validated with optogenetic experiments in behaving flies. This is not in direct tension with our trained-wiring result: their test is of structural sufficiency under randomly-parameterized simulation, not of trained real wiring against trained random wiring, the comparison this paper’s own finding turns on. But it is a useful reminder that the erosion of a real-versus-random signal under training, the central finding of this paper, is not evidence that wiring structure is uninformative in general; it may instead reflect something specific to task optimization eroding a signal that untrained or minimally-parameterized structure alone can still carry.

### What biological wiring contributes, and what remains unexplained

Even where the trained comparison converges, structure in the untrained comparison is informative about what the connectome contributes before training reshapes it. Real wiring’s own within-ensemble consistency is itself lower on OFF-only structure (*r* = 0.634) than ON-only structure (*r* = 0.838), an asymmetry that survives the within-polarity decomposition used to rule out the ON+OFF confound elsewhere in this paper, and for which we do not currently have a mechanistic account. Two further asymmetries compound this puzzle rather than resolve it. First, the null scheme that preserves *less* structure, Erdős–Rényi, consistently retains *more* residual similarity to real wiring, untrained, than the gentler degree-preserving swap does, the opposite of the naive expectation that a more aggressive scramble should erase more signal. Second, this same scheme, once trained, is the one that reaches full statistical indistinguishability from real wiring, and specifically on the same OFF-polarity structure where real wiring’s own internal consistency is weakest. Whether these three observations, the OFF-polarity consistency gap, Erdős–Rényi’s untrained residual similarity, and its trained OFF-polarity convergence, are causally connected or an accumulation of independent coincidences is a genuinely open question this paper raises but does not resolve.

### On measuring what cannot be measured

Two methodological failures, caught during this work rather than after publication, are worth reporting as general lessons rather than as footnotes. An early biological reference was 97.8% explained by circular stimulus structure; nothing in a standard RSA pipeline asks what a reference is capable of measuring before correlating something against it, and a plausible-looking, significant correlation was in fact almost entirely circular ordering. The fix, correlating the reference against a pure circular-distance model before trusting anything else about it, is one line of code and is now standard practice throughout this paper. A second failure was subtler and specific to this project’s own statistical choice, not to this particular numerical regime: ensemble-mean RDM correlation, computed by averaging every model’s response pattern before comparing, can understate the true real-versus-random gap so severely that, on one stimulus condition, it produced a number exceeding the within-real ceiling itself, a result an honest comparison cannot produce. Individual-pairwise comparison, which computes every real-versus-real and real-versus-null correlation independently before summarizing, does not have this failure mode, and we report it as primary throughout. Both failures share a structure: a plausible-looking statistic was computed and reported without first checking whether the instrument or the statistic itself was capable of the measurement being asked of it. We regard this checking discipline, not any single corrected number, as the more durable contribution of this paper’s methodology.

### Relationship to neural predictivity

Lappalainen et al. (Lappalainen et al., 2024) validated the Flyvis ensemble against recorded T4/T5 responses directly, showing that models with lower task error also predict single-neuron tuning more accurately. Our Method A result, once size-matched, is a genuine null on the same broad comparison at the population level: trained real and trained-random wiring are statistically indistinguishable against the same class of biological reference. This is not in tension with Lappalainen et al.’s finding, which compares real, trained connectomes against each other by task performance, not against trained-random wiring; the two results describe different comparisons. The RSA-based framework used throughout this paper complements single-neuron predictivity by operating at the population level without requiring single-unit recordings and without assuming a particular decoding scheme: predictivity asks whether a model’s responses match specific recorded neurons; representational geometry asks whether the relational structure of population responses is preserved, a question that remains answerable even where no single-neuron ground truth exists.

### Limitations and future directions

Several limitations qualify the current results. First, both biological references, Maisak-derived and Henning-derived, cover only T4/T5 (a handful of the 65 modeled cell types), and the Henning reference itself remains substantially circular (63.5–82.3% of variance) even after validation; a reference spanning more cell types with genuinely heterogeneous tuning would strengthen every biology-mediated comparison in this paper. Based on direct correspondence (2026), a non-circular, stimulus-matched physiological reference of this kind is being actively pursued by Jakob Macke’s lab, including PhD student Linda Ulmer, as part of a broader survey effort to align connectome-constrained models with neural activity measurements; we were not aware of an existing dataset meeting this need at the time of writing. Second, the Erdős–Rényi Method A result, though it does not survive size-matching as reported, is resampled from *n* = 10 down from *n* = 50, not scaled up; the resampled *p*-values (range 0.04–0.21) were consistent with either a genuinely null effect or a real but underpowered one. We have since trained an extended Erdős–Rényi ensemble (*n* = 24, one network excluded as unstable, a constant, non-direction-selective output) and evaluated it directly against real wiring (*n* = 50) without resampling: the difference remains non-significant (raw reference, *p* = 0.49), consistent with the resampled analysis’s null interpretation rather than an underpowered real effect. A related but distinct extension, to the negative-trending snapshot test of §3.4 (Table 10), is also now complete for both null schemes: degree-preserving swap reaches significance at *n* = 25 (*p* = 0.0146) where the original *n* = 10 sample did not, while Erdős–Rényi’s own snapshot result moves in the opposite direction, from 100% negative (*p* = 0.002) at *n* = 10 to 67% negative, not significant (*p* = 0.152), at *n* = 24. Read together, both extensions agree for Erdős–Rényi specifically: its original apparent negative signal does not survive a larger, independently-evaluated sample, on either test. Third, the one case where training closes the gap completely, Erdős–Rényi on OFF-polarity structure, does so only at the final training checkpoint, not gradually or early; whether this reflects a genuine, late-emerging effect or a boundary case that happened to land on one side of a significance threshold remains open, and we report it with that caveat rather than as a settled result. Fourth, the retraining-based instability test (§3.5) tests only eight seed networks; the permutation test’s own heterogeneity, some networks cluster tightly across retraining while others scatter or reverse entirely, means a null result on wiring-identity may reflect genuine training-process randomness or simply insufficient power at this sample size, and we do not attempt to distinguish these. Fifth, two individual networks in that same test, 0007 and 0009, show internal training dynamics, a mid-training reversal producing three distinct retrained outcomes from identical wiring, and a consistently-reproduced early-swing-then-recovery shape, respectively, that are not explained by the aggregate wiring-versus-noise result and that we have not pursued with the dynamical-systems tools such signatures would warrant. Sixth, this paper is scoped to a single species and a single connectome-constrained modeling framework; a parallel line of work applying the same representational-geometry approach to mouse visual cortex is reported separately. Finally, this paper does not address which specific subset of neurons or cell types would be minimally sufficient, in combination with connectome data, to constrain a model’s dynamics; recent theoretical work (Beiran and Litwin-Kumar, 2025) shows that connectome constraints alone often fail to constrain the dynamics of a task-performing network, and develops a concrete optimal-selection procedure, ranking neurons by how much their recording would reduce error in a model’s unconstrained parameter directions, that identifies which small subset of recordings is most informative and substantially outperforms random selection. Applying a comparable procedure to identify an optimal or near-minimal recording subset for our own system, rather than assuming, as we do throughout, that all 65 cell types are available, is an open direction this paper does not pursue.

### Preliminary explorations prompted by external review

Following circulation of an earlier draft, two questions raised in correspondence prompted three follow-up analyses, summarized in Table 12. None of these has undergone the validation applied to other findings in this paper: each is a single run on a single stimulus condition (or, for the cross-individual test, the one dataset available), without permutation testing, replication across conditions, or comparison against a proper null distribution. They are reported here only as preliminary indications that these directions are tractable and their headline numbers, not as validated findings on par with the rest of this paper.

**Table 12:** Preliminary follow-up analyses, not validated to the standard of the rest of this paper. All results single-run, ON-only condition where applicable.

| Question | Method | Result |
| --- | --- | --- |
| Which cell types are most informative for reconstructing the trained population’s representational geometry? | Greedy selection maximizing reconstruction of the full-population RDM | 4 of 65 cell types reach Spearman $r \geq 0.8$; dominated by T4 subtypes |
| Does connectome structure alone (no activity) predict this same ranking? | In-/out-degree, computed directly from the connectome, compared against the activity-based ranking above | In-degree $\rho = 0.41\text{--}0.51$ ($p < 0.001$); out-degree $\rho = 0.23$ ($p = 0.07$, n.s.); T4 subtypes not among highest-degree cell types |
| Can cross-individual (donor) recordings substitute for within-individual recordings in the Henning reference? | Leave-one-fly-out (14 flies) vs. matched random split-half, recomputed on the same raw data | Leave-one-out $\rho = 0.81$ (raw) / $0.77$ (von Mises) vs. split-half $\rho = 0.99 / 0.98$; gap of $0.18\text{--}0.22$; per-fly $\rho$ ranged $0.47\text{--}0.94$ |

The first two rows were independently reproduced on a second machine with closely matching numbers; the third was reproduced with matching results to three decimal places. Read together, they suggest three things: this class of follow-up question is empirically tractable with the existing pipeline: connectome degree alone is a weak proxy for which units matter most; and the pooled, cross-individual biological reference used throughout this paper substitutes for within-individual data meaningfully, but not perfectly, a real, quantified caveat on the biology-mediated comparisons in §3.2 and §3.3, not merely a theoretical one.

## 5 Conclusion

This paper establishes representational geometry as a genuine, population-level fidelity signal for connectome-constrained models: real wiring is reliably distinguishable from random wiring before training, corroborated independently by CKA, but that distinction narrows substantially and in one specific case disappears entirely once both are trained to task adequacy, the direct test Brunton et al.’s finding raises but does not itself perform. Reaching this result required first diagnosing and repairing the biological reference used to validate it, the original reference was measuring circular stimulus structure more than direction tuning, and correcting a separate statistical artifact in how representational similarity itself was measured. We report this as an honest, largely null result on the training question, not the confirmation we set out to find, alongside a validated non-circular biological reference, a diagnosed methodological pitfall, and a set of open, quantified limitations, resources we intend as useful starting points for evaluating fidelity in connectome-scale emulations more broadly.

Beyond the specific findings above, we see this work as useful in three ways. For researchers building connectome-constrained models of other systems, the individual-pairwise RSA methodology, the diagnosed ensemble-mean statistical artifact, and the circularity check for biological references are directly transferable tools, not specific to the fly visual system, and worth applying before trusting a fidelity claim built on similar comparisons. For the growing whole-brain-emulation community, this paper offers a concrete, empirically tested instance of the kind of fidelity metric that broader conceptual frameworks have called for but not yet validated against real data (Linssen and Koene, 2025). And for anyone tempted to treat a model’s ability to reproduce behavior as sufficient evidence of biological fidelity, our results, particularly the case where training closes the real-versus-random gap completely, are a direct, quantified caution: behavioral adequacy and structural fidelity are not the same claim, and only one of them is guaranteed by matching output alone.

## Data and Code Availability

All code, experiment scripts, saved results (.npz files), and figures are available at https://github.com/michaela10c/connectome-fidelity.

The trained-random training and evaluation pipeline (production.py, randomize_connectome_schemes.py), the Henning reference construction (build_henning_reference.py, build_reference_from_raw.py), and the full-checkpoint trajectory evaluation (evaluate_full_trajectory_all_conditions.py, analyze_full_trajectory.py) are included in the repository, along with the individual-pairwise RSA scripts (test_item1_all_null_schemes.py, analyze_within_polarity_decomposition.py) used throughout §3.1 and §3.3.

The pretrained Flyvis ensemble (Lappalainen et al., 2024) is available at https://github.com/TuragaLab/flyvis.

A parallel line of work applying this same representational-geometry framework to mouse visual cortex (MICrONS) is available separately at https://github.com/michaela10c/connectome-fidelity-microns; it is supplemental to this paper, not part of its reproduction pipeline or its reported results.

## Acknowledgments and Disclosure of Funding

Michael G. Zhou is supported by the Georgia Tech ECE Fellowship and a Graduate Teaching Assistantship. The authors thank the Flyvis team (Lappalainen et al.) for releasing the pretrained Drosophila visual system ensemble as an open-source resource.

This paper’s text was drafted with substantial assistance from Claude (Anthropic), including full drafts of the Methods, Results, and Discussion sections, built directly from the author’s own experimental scripts, saved results, and verified source citations. All experiments, analyses, and scientific conclusions are the author’s own; AI-assisted text was checked against primary sources throughout, and corrections were made where discrepancies were found.

